# NextPolish2:a repeat-aware polishing tool for genomes assembled using HiFi long reads

**DOI:** 10.1101/2023.04.26.538352

**Authors:** Jiang Hu, Zhuo Wang, Fan Liang, Shanlin Liu, Kai Ye, De-Peng Wang

## Abstract

The high-fidelity (HiFi) long-read sequencing technology developed by PacBio has greatly improved the base-level accuracy of genome assemblies, but these assemblies still contain some base-level errors, particularly within the error-prone regions of HiFi long reads. However, existing genome polishing tools usually introduce overcorrections and haplotype switch errors when correcting errors in genomes assembled from HiFi long reads. Here we describe an upgraded genome polishing tool - NextPolish2, which can fix base errors remaining in those “highly accurate” genomes assembled from HiFi long reads without introducing excess overcorrections and haplotype switch errors. We believe NextPolish2 has a great significance to further improve the accuracy of Telomere-to-Telomere (T2T) genomes. NextPolish2 is freely available at https://github.com/Nextomics/NextPolish2.

## Introduction

Complete and accurate genomes provide fundamental tools for scientists to capture a full spectrum of the genomic variants and use that information to understand the evolutionary basis of various diseases and other biological phenotypes [1]. Hence complete and gapless genome, also known as T2T genome, has been emerging as a new hotspot in the field of genomics [2–7]. Typically, we obtain a T2T genome with datasets including both high-accuracy PacBio HiFi long reads and Oxford Nanopore Technologies (ONT) error-prone ultra-long reads [2]. Compared to those genomes that were generated using noisy long reads, genomes obtained using HiFi long reads have considerably higher qualities - much less errors at the level of single nucleotides and small insertions and deletions [8,9]. However, they still contain a handful of assembly errors in chromosomal regions where HiFi long reads stumble as well, such as homopolymer or low-complexity microsatellite regions (**Figures S1 and S2**). Additionally, a typical gap-filling step is accomplished using ONT ultra long reads which contain a certain amount of errors that need to be corrected [10]. Hence, the current T2T genomes assembled using the cutting-edge sequencing platforms still require further improvement in terms of consensus accuracy. For example, the human genome T2T consortium has applied multiple tools and extensive manual validation to increase the assembly quality value (QV) from 70.2 to 73.9 for the T2T assembly of a human genome (CHM13) [10].

Error correction for a T2T genome assembly is challenging because (i) complex segmental duplications and large tandem repeats, such as centromeric satellite arrays, could potentially induce overcorrections or false-negative corrections; (ii) local haplotype needed to be maintained; and (iii) technology-specific biases of different sequencing platforms [11]. Therefore, although there are many state-of-the-art polishing tools available, such as Pilon [12], Racon [13] and NextPolish1 [14] et. al. They were designed for error correction of genomes assembled from noisy long reads and can hardly handle T2T genome assemblies.

Here, we present an upgraded genome polishing tool, NextPolish2, for error correction of T2T genomes constructed mainly using HiFi long reads. Compared to the up-to-date polishing pipeline (Racon + Merfin [10], hereafter referred to as RM) adopted to polish the human T2T genome assembly of CHM13, NextPolish2 can fix base errors in “highly accurate” draft assemblies without introducing overcorrections, even in regions with highly repetitive elements. Through the built-in phasing module, it can not only correct the error bases, but also maintain the original haplotype consistency. In fact, our evaluation shows it even slightly reduce switch errors in heterozygous regions.

## Algorithm

NextPolish2 follows the Kmer Score Chain (KSC) algorithm of its previous version to perform an initial rough correction [14], and detect low-quality positions (LQPs) where the chosen alleles account for ≤ 0.95 of the total during a traceback procedure (**Figure 1A**). Next, it merges the adjacent LQPs into low-quality regions (LQRs), and then for each LQRs it extracts kmers from the HiFi long reads that can map across those LQRs. The kmer set of each LQR is subsequently filtered using kmer datasets generated from high quality short reads (**Figure 1B**). After that, it defines kmer sets that contain ≥ 2 valid kmers as heterozygous and uses them to calculate weights between reads spanning the same LQRs. And then it applies the Louvain community detection algorithm [15] to group reads from the same haplotype or repeat copy. We define two conflict communities (*C*_*1*_, *C*_*2*_) as *weight(C*_*1*_, *C*_*2*_) < *0*, which are located in the same region but from different haplotypes or repeat copies (**Figures 1C and 1D**). For the conflict communities, we only use the community that contains the most reads or shares the most kmers with the reference sequence based on user settings, and remove reads from other communities (**Figure 1E**). We repeat the above procedure until all conflict communities are resolved (the number of iterations can be adjusted according to user settings, **Figure 1F**), and then use the KSC algorithm to generate a draft consensus sequence. The draft consensus sequences may still contain a small number of LQRs. For those LQRs not spanned by any valid kmers, we use the kmer from the reference sequence as the correct kmer to avoid overcorrection. For LQRs spanned by multiple valid kmers, the kmer with a highest number is defined as the correct one. Finally, we update the draft consensus sequence using these correct kmers and generate the final consensus sequence.

**Fig. 1:**
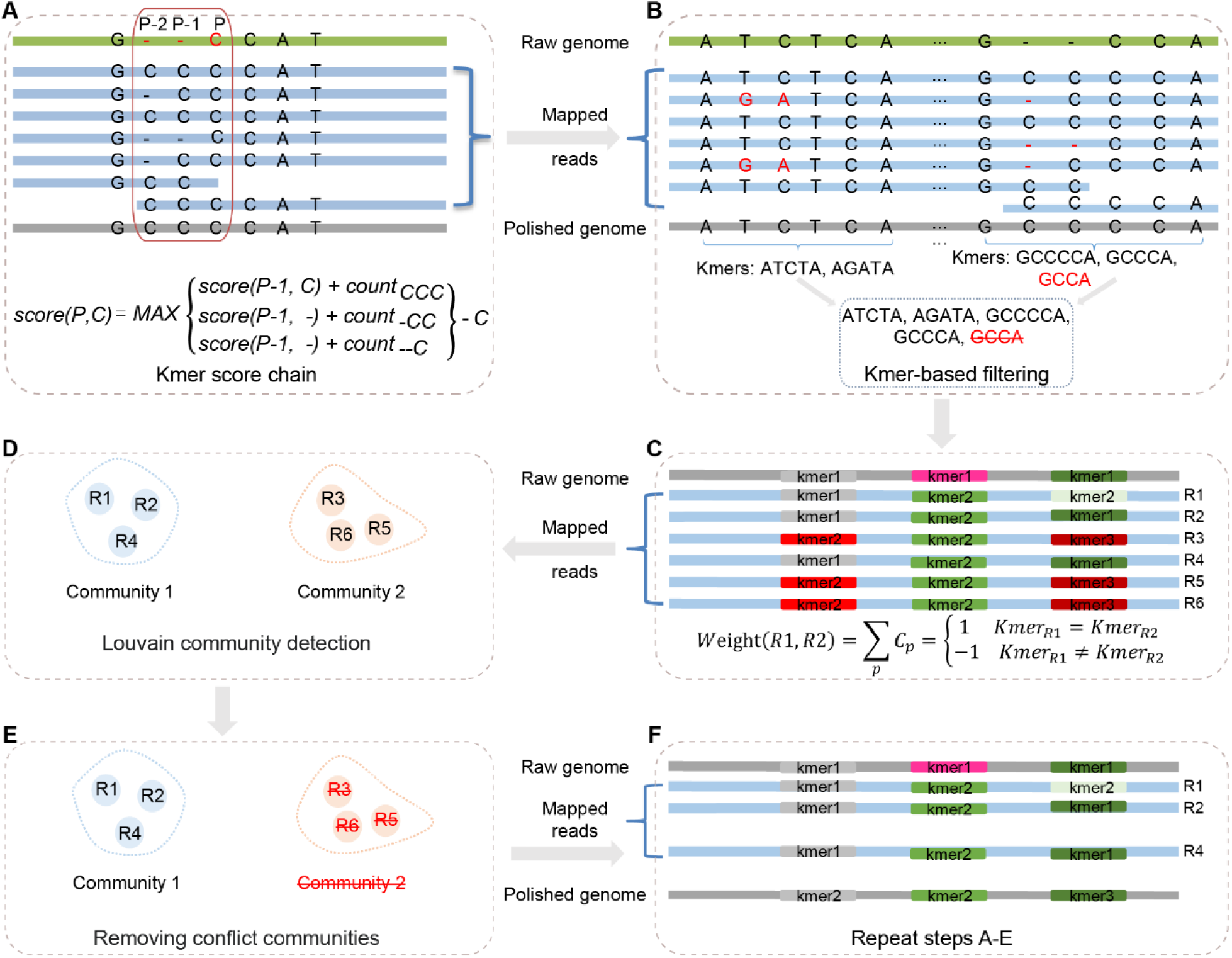
NextPolish2 pipeline. (**A**) The schematic of the K-mer score chain algorithm. The score of the base ‘C’ at position P is the maximum score of its predecessor bases (C/deletion) at position P-1, plus the count of their corresponding 3-mers (CCC, -CC and --C), and then minus the valid depth (6) of position P; (**B**) K-mers at LQRs are extracted and filtered using the kmer datasets. (**C**) Weights between reads are calculated using the count of kmers. (**D**) Reads are grouped using the Louvain community detection algorithm. (**E**) Only use one community for multiple conflicting communities, and discard reads in the communities to be removed. (**F**) Repeat steps A-E until there are no conflict communities.

## Performance

We evaluated the performance of NextPolish2 against RM using three datasets, including (i) HiFi long reads and Illumina short reads simulated based on a simulated highly heterozygous diploid *Arabidopsis thaliana* genome; (ii) published sequencing data of *A. thaliana* (Col-XJTU) and (iii) *Homo sapiens* (HG002, **Table S1**). We first applied Hifiasm (v0.18.5) [16] to obtain genome assembly for each dataset. In addition to a primary assembly (a complete assembly with long stretches of phased blocks), we generated two haplotype-resolved assemblies (two complete assemblies consisting of haplotigs, representing an entire diploid genome) for the human genome with the available trio binning dataset. All the assemblies reached QV scores of ca. 48∼60 (**Table 1**). Then, we mapped HiFi long reads of each sample onto their corresponding genome assemblies using Winnowmap2 (v2.03) [17], a repeat-aware alignment tool that was adopted in the RM polishing pipeline. After that, we applied both Nextpolish2 and RM pipeline to conduct error correction separately for each genome assembly. Finally, we applied yak (v0.1, for the simulated datasets, https://github.com/lh3/yak) and merqury (v1.3, for the actual biological datasets) [18] to evaluate QV and switch errors of the reference genomes and the polished genomes using kmers from Illumina reads. In addition, we applied meryl (v1.3, https://github.com/marbl/meryl) to detect kmer changes between the reference genomes and the polished genomes to evaluate the challenge of overcorrection.

**Table 1:** Statistics of genome polishing results.

| Source | Software | QV | Switch error rate (‰) | Changed kmers | Potential overcorrection kmers | Wall clock time<sup>b</sup> (minute) |
| --- | --- | --- | --- | --- | --- | --- |
| *A. thaliana*<br>(simulated data, primary contigs) | Hifiasm (primary) | 47.67 | 1.99 |  |  |  |
|  | Racon | 43.12 | 765.95 | 6,384,788 | 89,878 | 20.77 |
|  | NextPolish1 | 45.01 | 1058.34 | 6,159,005 | 36,758 | **2.00** |
|  | Racon + Merfin | 52.18 | 737.45 | 6,207,453 | 114 | 26.73 |
|  | NextPolish2 | **65.42** | **0.35** | **25,869** | **0** | 2.53 |
| *A. thaliana*<sup>a</sup><br>(Col-XJTU, primary contigs) | Hifiasm (primary) | 58.03 |  |  |  |  |
|  | Racon | 46.58 |  | 53,462 | 213,624 | 14.33 |
|  | NextPolish1 | 57.21 |  | 69,606 | 3,134 | **1.07** |
|  | Racon + Merfin | 63.89 |  | 7,220 | 48 | 13.88 |
|  | NextPolish2 | **64.26** |  | **1,477** | **0** | 6.28 |
| *H. sapiens*<br>(HG002, primary contigs) | Hifiasm (primary) | 60.25 | 0.15 |  |  |  |
|  | Racon + Merfin | **63.52** | 5.40 | 17,835,299 | 2,895 | 544.64 |
|  | NextPolish2 | 62.87 | **0.14** | **32,508** | **45** | **88.72** |
| *H. sapiens*<br>(HG002, paternal contigs) | Hifiasm (trio) | 59.77 | 0.21 |  |  |  |
|  | Racon + Merfin | 63.44 | 0.69 | 3,711,007 | 1,889 | 297.92 |
|  | NextPolish2 | **63.49** | **0.20** | **588,415** | **316** | **92.65** |
| *H. sapiens*<br>(HG002, maternal contigs) | Hifiasm (trio) | 59.78 | 0.33 |  |  |  |
|  | Racon + Merfin | 63.23 | 1.60 | 4,940,002 | 2,088 | 320.21 |
|  | NextPolish2 | **63.29** | **0.30** | **403,062** | **183** | **109.05** |

a: Unable to evaluate switch error rate due to missing parental sequencing dataset.
b: Only the time for genome correction is counted, and the time for reads mapping is not included.
Hifiasm (primary): primary hifiasm assembly. Hifiasm (trio): haplotype-resolved hifiasm assembly with trio binning. The best value for each metrics is indicated with bold type. All the software were tested on the same computer with 5 CPUs and 128 GB RAM of memory.

### Correction accuracy

*Regarding* the *A. thaliana* genome, NextPolish2 outperformed RM for both the simulated and actual biological datasets. NextPolish2 corrected more errors and thus resulted in higher QVs. For the human genome, the performance of the two analysis pipelines was evenly matched when worked on the haplotype-resolved assemblies, while the RM pipeline generated a polished genome with higher QV than that of NextPolish2 when worked on the primary assembly. However, it is worth noting that the QV advantage of RM pipeline came at the cost of breaking haplotype blocks and introducing more haplotype switch errors. It shows that the polished assemblies of RM contained more switch errors than that of NextPolish2 for all the test datasets (Table 1). Given that the RM pipeline was developed to correct the CHM13 genome assembly that is a homozygous cell line based and thus contains a limited number of heterozygous loci, it may not design any particular modules to deal with haplotype switch errors.

### Overcorrections

We calculated two metrics: “changed kmers” and “potential overcorrection kmers” to evaluate the overcorrection issue, of which the former is the count of kmers that present in a polished assembly but not in its reference genome, and the latter is the count of kmers that present in a polished assembly but neither in its reference genome nor in Illumina short reads of the same sample. If a polishing tool introduces too many overcorrections, the polished assembly will contain lots of “changed kmers” and “potential overcorrection kmers”, because introducing a new kmer with length of *k* may indirectly introduce ≤ 2 × (*k* − 1) kmers overlapping with this new kmer. Compared to NextPolish2, RM introduced ∼6.31 – 548.64 times more “changed kmers” without significant improvement of QV, which means that RM overcorrected many authentic sequences of bases on the reference genomes. The fact that RM introduced 5.98-64.33 times more “potential overcorrection kmers” than NextPolish2 on human genome assemblies also told the same thing (**Table 1**).

To verify the overcorrection estimation, we identified 158 transposable elements (TEs) in the simulated *A. thaliana* genome and used them to evaluate the error correction performance of the polishing tools for those highly repetitive regions on the genome. By comparing the mapping identity rate between the assembled genomes and the reference, we found that a total of 149 TEs were successfully assembled, but only 68.46% of them can map to their corresponding TE references with an identity rate of 100%. After genome polishing, NextPolish2 increased the ratio from 68.46% to 91.95% and no TE was introduced overcorrections after genome polishing, but RM decreased the ratio from 68.46% to 12.75%, and about 80.54% TEs had a lower identity after genome polishing (**Table S2**).

### Computational resource consumption

On running time, NextPolish2 accomplished error corrections considerably faster than RM ∼2-6 times and ∼11 times faster for the real-world and simulated datasets, respectively (Table 1). The further improvement of the simulated dataset could be attributed to the heterozygous issue mentioned above.

Additionally, we found that the genome polishing tools designed for long noisy reads, such as Racon and NextPolish1, introduced more errors than what they corrected (**Table 1**) and thus are not recommended for error correction of genomes assembled using HiFi long reads.

## Discussion

NextPolish2 is a fast open-source polishing tool specifically developed for error correction of genomes assembled from HiFi long reads. It is an upgraded version of NextPolish1 and can also work on genomes assembled from noisy long reads, as well as those gap regions that are filled with sequences generated from ONT ultra long reads in T2T genomes.

We found the polished genomes still contain some errors, of which most of their corresponding genomic regions were not covered by Illumina short reads or demonstrated high inconsistencies among the mapped short reads, which impedes the performance of NextPolish2 as it relies heavily on short reads to check whether a kmer contains errors. Therefore, we encourage users to use PCR-Free libraries and high-coverage short reads to minimize uncorrected errors caused by biases inherent in short read sequencing technologies, especially for T2T genome projects that pursue extremely-high-quality genome assemblies.

## Code availability

NextPolish2 is implemented in Rust. The source code as well as results of the benchmark tests are freely available from https://github.com/Nextomics/NextPolish2

## Supplementary Figures

**Supplementary Figure S1.**
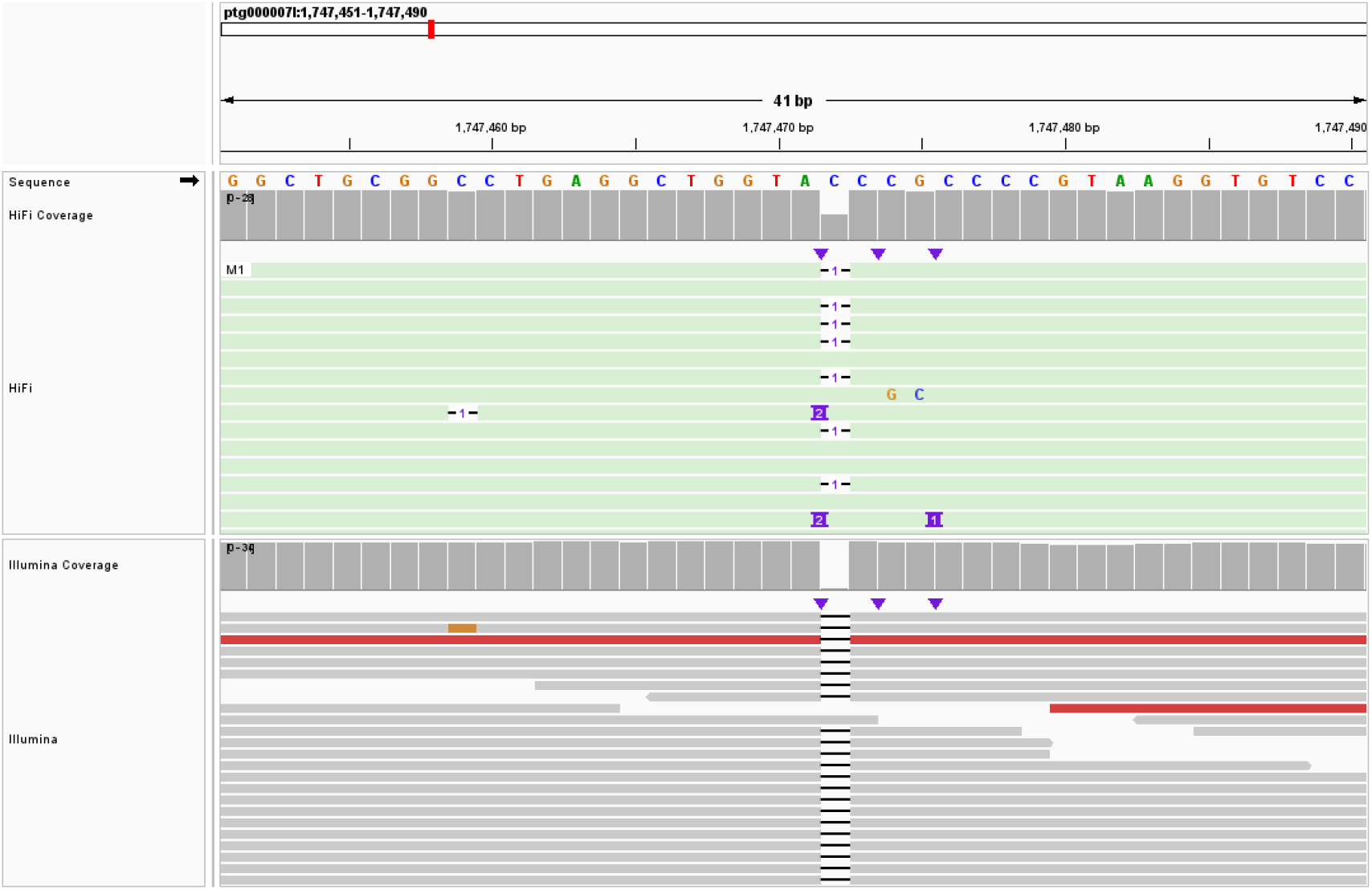
Integrative genomics viewer screenshot of a deletion error in the reference genome.

**Supplementary Figure S2.**
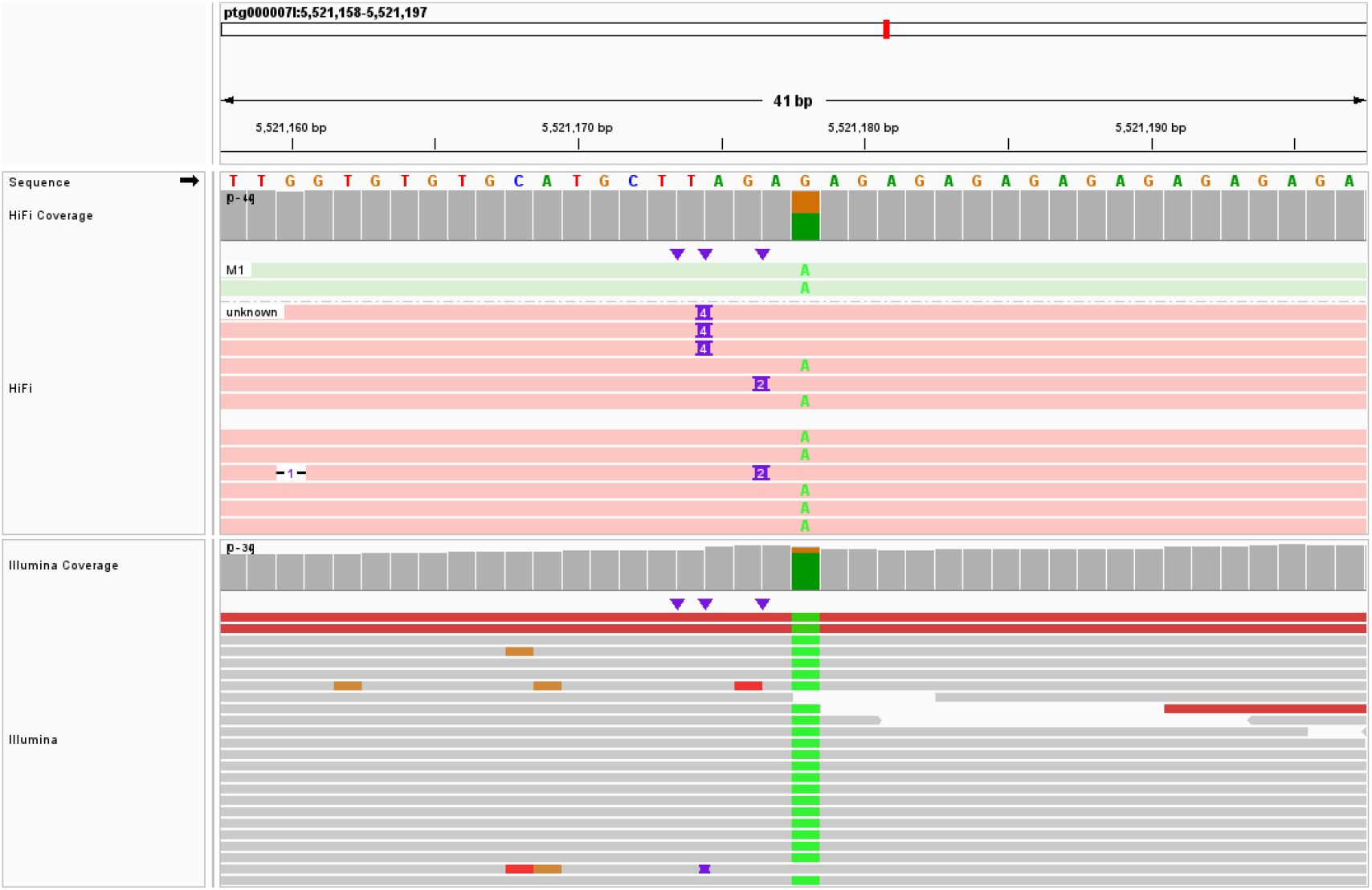
Integrative genomics viewer screenshot of a SNV error in the reference genome.

## Supplementary Tables

**Supplementary Table S1.** Statistical information of the datasets used in this study.

| Source | Read type | Bases (bp) | Average Reads Length (bp) | Base coverage |
| --- | --- | --- | --- | --- |
| *A. thaliana*<br>(simulated data) | Illumina | 13,207,830,000 | 150 | 100.00 |
|  | HiFi | 8,343,317,687 | 12,956 | 63.17 |
| *A. thaliana* | Illumina | 13,696,431,300 | 150 | 104.82 |
|  | HiFi | 4,696,718,342 | 15,098 | 35.95 |
| *H. sapiens*<br>(HG002) | Illumina | 100,892,786,960 | 148 | 32.75 |
|  | HiFi | 110,549,151,396 | 14,971 | 35.88 |
| *H. sapiens* (HG003) | Illumina | 101,741,039,192 | 148 | 33.02 |
| *H. sapiens* (HG004) | Illumina | 101,309,565,616 | 148 | 32.88 |

**Supplementary Table S2.** Accuracy of transposable elements in pre- and post-polishing assemblies of the simulated *A. thaliana* genome.

| Source | Software | A total of 149 transposable elements |  |
| --- | --- | --- | --- |
|  |  | 100% mapping identity (%) | Lower identity after polishing (%) |
| *A. thaliana*<br>(simulated data, primary contigs) | Hifiasm (primary) | 68.46 |  |
|  | Racon + Merfin | 12.75 | 80.54 |
|  | NextPolish2 | 91.95 | 0.00 |

Hifiasm (primary): primary hifiasm assembly. The identity was defined by minimap2 and only the primary alignments of each transposable element were used for evaluation.

## Notes

https://github.com/Nextomics/NextPolish2

### Competing Interest Statement

The authors have declared no competing interest.

